# Immune pathways and perinatal environmental exposures contribute to epigenetic gestational age prediction and acceleration

**DOI:** 10.1101/2025.06.13.659090

**Authors:** Amy A. Eapen, Ian M. Loveless, Mingming Pan, Xiaoyu Liang, Audrey Urquhart, Jennifer Straughen, Andrea E. Cassidy-Bushrow, Alexandra R. Sitarik, Neil Simmerman, Emma E. Thompson, Leah Kottyan, Carole Ober, Christine C. Johnson, Edward Zoratti, Albert M. Levin

**Affiliations:** Division of Allergy and Clinical Immunology, Department of Internal Medicine, Henry Ford Health + Michigan State University Health Sciences, Detroit, MI; Department of Medicine, College of Human Medicine, Michigan State University; Center for Bioinformatics, Henry Ford Health + Michigan State University Health Sciences, Detroit, MI; Department of Public Health Sciences, Henry Ford Health + Michigan State University Health Sciences, Detroit, MI; Department of Epidemiology and Biostatistics, Michigan State University, Lansing, MI; Department of Obstetrics, Gynecology and Reproductive Biology, College of Human Medicine, Michigan State University, East Lansing, MI; Department of Pediatrics and Human Development, College of Human Medicine, Michigan State University, East Lansing, MI; Department of Women’s Health Services, Henry Ford Health + Michigan State University Health Sciences, Detroit, MI; Department of Human Genetics, University of Chicago, Chicago, IL; Center of Autoimmune Genomics and Etiology, Division of Eosinophilic Disorders, Cincinnati Children’s Hospital Medical Center, Cincinnati, OH

**Keywords:** Gestational age, Epigenetic gestational age clock, Prenatal environment

## Abstract

DNA methylation (DNAm), capturing biological gestational age (GA) and epigenetic gestational age acceleration (EGAA), can be modified by environmental exposures. The Asthma&Allergy array is a new DNAm array developed with content focused on asthma and allergy loci. The association between content on the Asthma&Allergy array and chronological GA and EGAA has not been evaluated alone or in the context of perinatal exposures. We performed an epigenome wide association study(EWAS) based on chronological GA at single CpG sites and regions. We further constructed a multi-CpG site methylation model to predict chronological GA in cord blood from 391 newborn children from a Detroit-based birth cohort. Associations between perinatal environmental factors with GA, epigenetic gestational age (EGA), and EGAA were assessed. We identified 2,435 CpG sites associated with chronological GA. HLA class II (*HLA-DRB1,HLA-DQB1,HLA-DRB6*) were the most significantly associated with chronological GA. Our multi-CpG site model attained predictive accuracy (cross-validated Pearson’s correlation=0.75) comparable to other EGA methods. Using genes implicated in region-based analyses (n=395 regions), the pathways most significantly enriched with chronological GA-associated CpGs included T helper 1(Th1) and 2(Th2) activation, macrophage classical activation, and IL10 signaling, which were also enriched in at least one of the other published epigenetic clocks. In multi-exposure models, prenatal indoor pet exposure and unplanned C-section were associated with EGA deceleration, while infant’s first-born status was associated with EGAA. Our findings highlight enrichment for T cell modulated pathways and antigen presentation as biological processes enriched in chronological GA, as well as novel perinatal factors that may impact EGAA.

## Introduction

Chronological gestational age (GA) is associated with developmental maturity, and both pre– and post-term births are associated with increased risk of adverse outcomes in both the perinatal period and also later on in life^1–3^. However, clinically measured chronological GA (by last menstrual period or ultrasound) is only one determinant of GA and is influenced by genetic and environmental factors^4^. DNA methylation (DNAm), which is influenced by the environment and genetics, is a marker of biological GA, and has been employed to create epigenetic gestational age (EGA) clocks^5^. The discordance between one’s epigenetic age and chronological age is a measure of whether individuals are biologically aging faster or slower relative to their chronological age^6^, which has been termed epigenetic age acceleration, or epigenetic gestational age acceleration (EGAA) when examing GA. EGAA has been associated with risk of a variety of childhood-onset conditions including, but not limited to, developmental delay and cardiometabolic disorders^7,8^.

Existing DNAm-based EGA clocks have generally utilized CpG sites included in Illumina methylation arrays (HumanMethylation 27 K, 450 K, and HumanMethylationEPIC 850K (EPIC)). While these arrays provide differing degrees of epigenome-wide coverage and have been successful at developing accurate GA clocks, they comprise <5% of the CpG sites across the human epigenome. As such, they have not comprehensively probed the contribution of DNAm to GA in loci that are associated with conditions that are impacted by biological pathways in aging.

Recently, leveraging whole genome bisulfite sequencing (WGBS) and *in silico* evidence of gene regulatory regions^9^, Morin et al developed a custom-content DNAm array^10^ that includes enhanced coverage of CpGs overlapping with predicted enhancers and transcription factor bindings sites within loci associated with asthma and allergic disorders compared to the Illumina EPIC array: the Asthma & Allergy array. It covers CpGs that are not present in the Illumina EPIC array. As biological aging has shown to be associated with risk of these conditions and immune pathways, it would be enlightening to characterize DNAm captured on this array and GA^11^.

In the current study, we applied the Asthma&Allergy array to cord blood DNA from a large metro-Detroit birth cohort that is diverse in terms of socioeconomic status (SES), parental-reported race, and urban/suburban residence to investigate the association between chronological GA and DNAm. We first identified individual CpG sites and regions of DNAm associated with GA and determined whether these sites were associated with specific immune pathways. Second, we developed a DNAm GA clock from the Asthma&Allergy array and evaluated its accuracy in predicting chronological GA compared to previously published clocks. Finally, as the environment (e.g. environmental tobacco smoke exposure (ETS), pet exposure, pollutant exposure, stress)^12^ can modify DNAm, we also investigated the associations of prenatal and perinatal environmental factors with EGA and EGAA in both individual and multi-exposure models.

## Materials and Methods

### Study cohort

The Wayne County Health, Environment, Allergy and Asthma Longitudinal Study (WHEALS) is a birth cohort from southeastern Michigan that has previously been described^13,14^. Briefly, women were eligible for inclusion in the study if they were in their second or third trimester of pregnancy, were between 21 and 49 years of age, and lived in a predefined cluster of zip codes in Detroit and its surrounding Wayne County suburbs. A total of 1,258 pregnant women were enrolled from September 2003 to December 2007 with no exclusion based on GA. Informed written consent was obtained for cord blood collection, as well as the use of genetic data for future assays. Mothers had the option to opt out of the genetic portion if desired and still remain in the study. This research was approved by the Henry Ford Health Institutional Review Board, protocol approval number14914 and 1881-29 (genetics sub-consent approved September 28, 2004), and adhered to the Declaration of Helsinki guidelines.

Cord blood was collected at delivery, and genomic DNA was isolated from whole cord blood using the QIAGEN FlexiGene DNA Kit (Germantown, MD). Of the 1,258 recruited women, 763 children (76.1%) either completed a 2-year follow-up visit in the clinic or had blood drawn for measurement of immunoglobulin E (primary outcome of parent study). From this subset who also had cord DNA stored, we performed a random selection of 391 participants and assessed for DNAm using the Asthma&Allergy array. Gestational age was chart abstracted from the participant’s obstetrician’s calculation in the electronic medical record, either ultrasound-based or from last menstrual period.

### DNAm quality control and quantification

Raw methylation Illumina *idat* files from the 391 cord DNA Asthma&Allergy array were loaded into R version 4.2.1 using the *minfi* package^15^. A total of 45,954 CpG sites were included in the array before QC. Probes with detection p-values greater than 0.05, with single nucleotide polymorphisms SNPs at either the 3’ or 5’ locations, on the sex chromosomes, or capturing CpG sites not intended to be targeted by the array were removed, resulting in a remaining 45,296 CpG sites. Remaining probes were then quantile normalized using the *ENmix* package in R^18^. Lastly, the resultant CpG site beta values were converted to logit transformed M-values, which were used for subsequent analyses.

### Statistical methods

Descriptive statistics of demographic characteristics, pregnancy and delivery factors, and prenatal environmental exposures are summarized in Table 1. We used mean and standard deviation for quantitative measures and counts and percentages for categorical measures. Univariate association between each one of these factors and chronological GA was calculated using linear regression, with an F-test used to assess statistical significance.

**Table 1.**
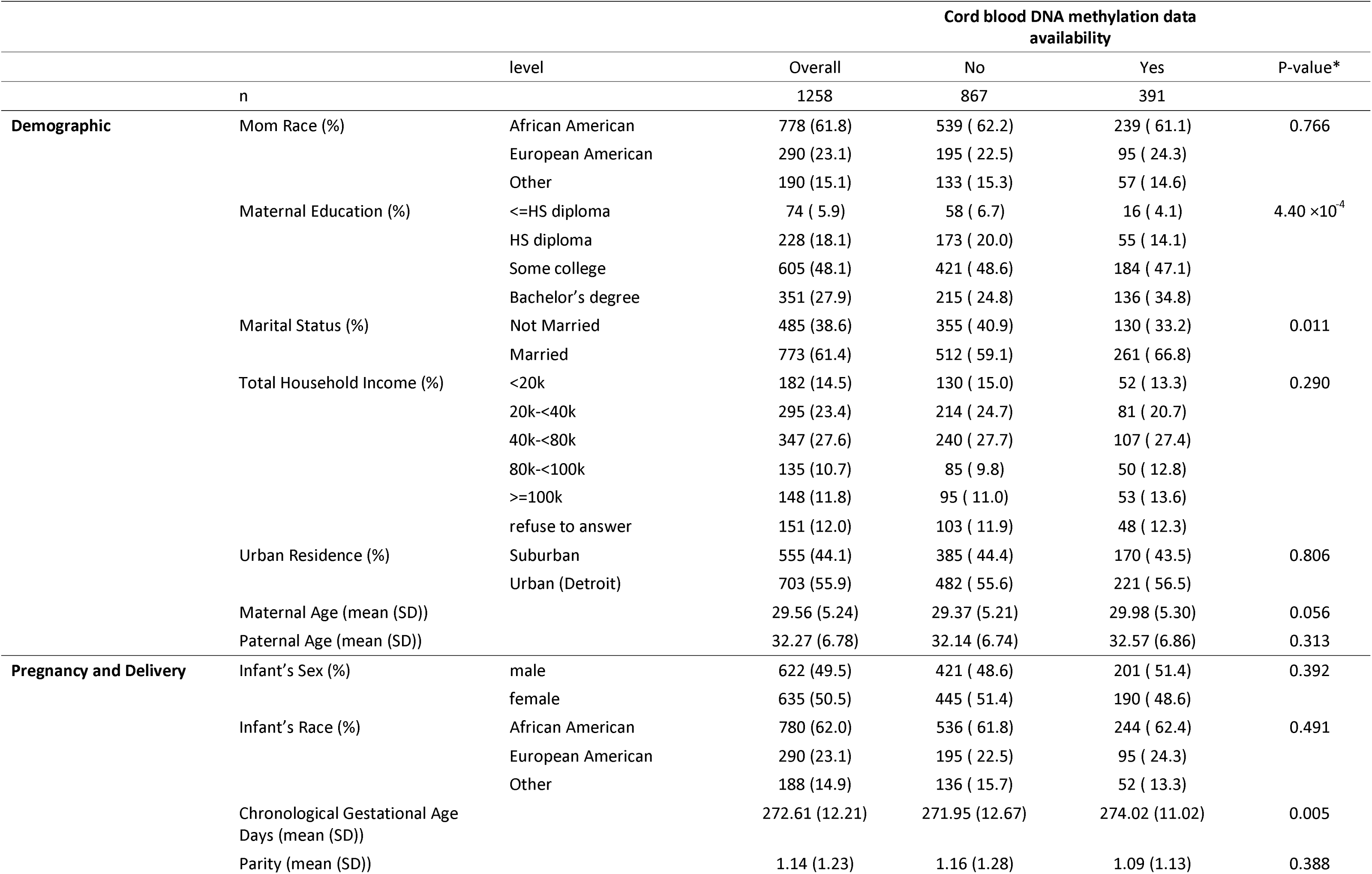
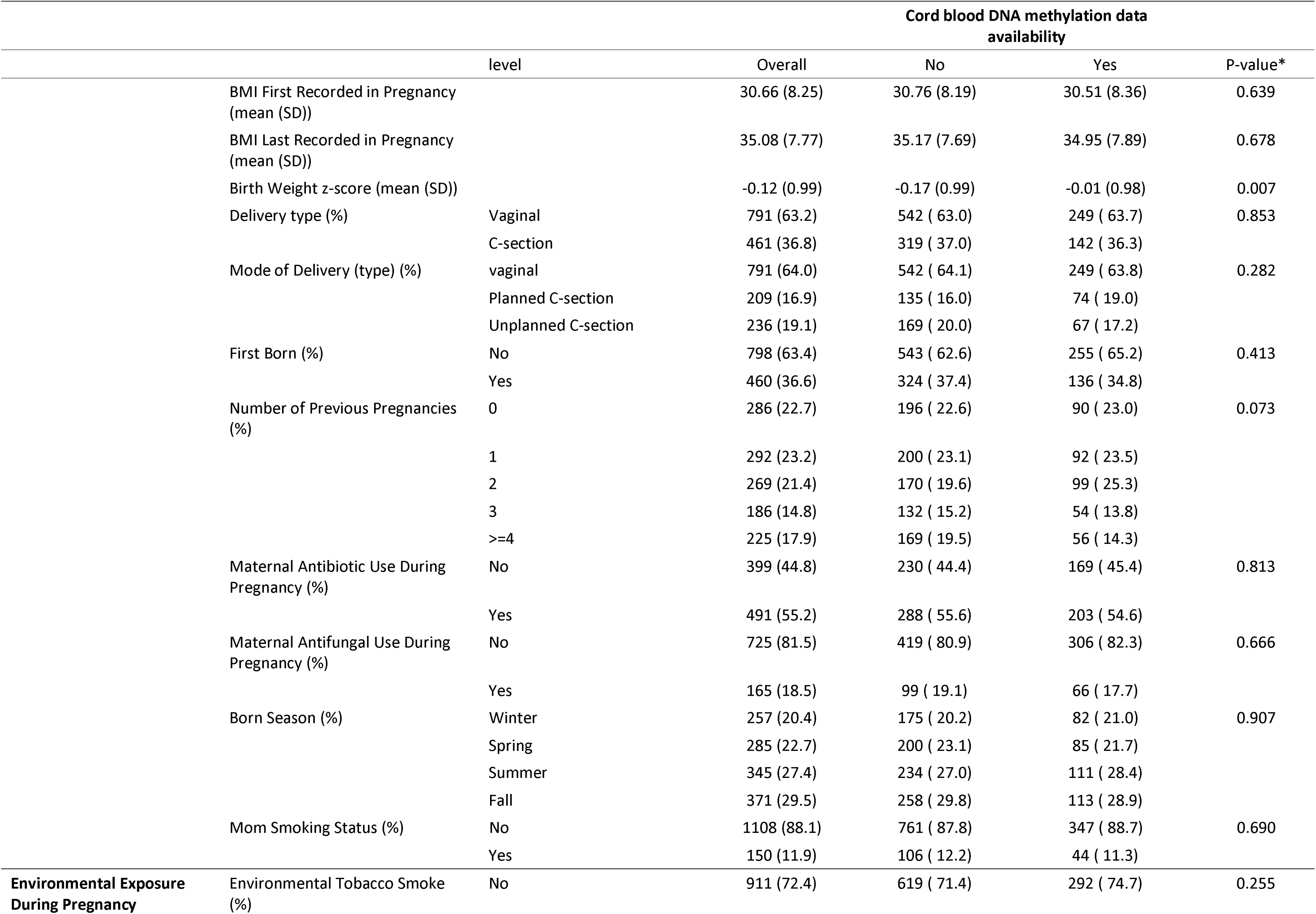
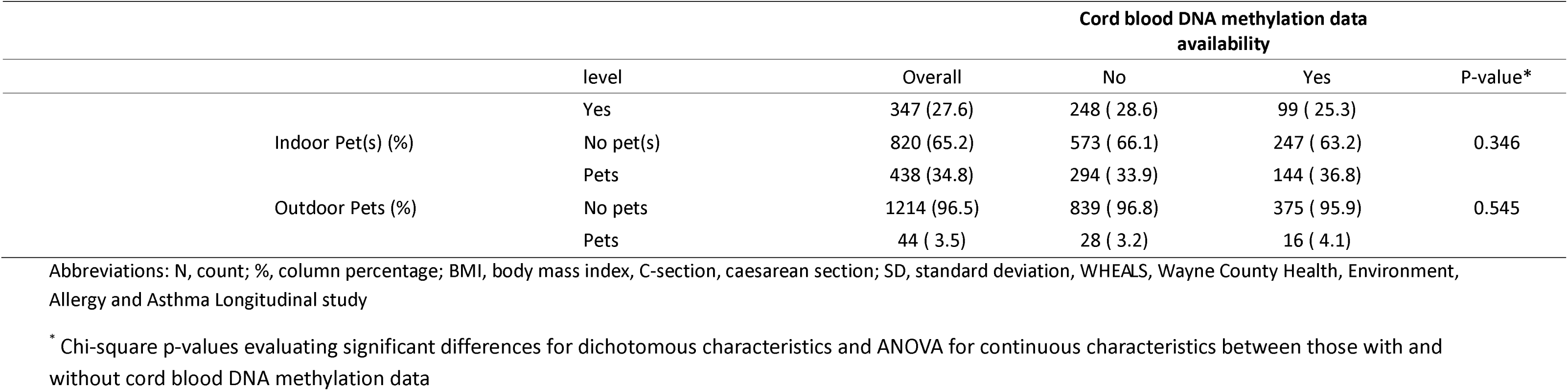
Demographic characteristics of WHEALS participants stratified by cord DNA methylation data availability.

First, an epigenome-wide association study (EWAS) was performed to identify CpG sites associated with chronological GA in days, adjusting for infant’s sex, parental-reported infant race, and latent features identified using the CorrConf package^16^. These latent features were identified after adjusting for infant’s sex, infant’s race, and GA in days. Association p-values were corrected for multiple-testing using an FDR threshold of 0.05^17^. For the region-based assessment, we identified differentially methylated regions (DMRs) based on combinations of p-values from the single site EWAS using *comb-p*^18^, which was implemented in the Enmix R package^19^. Individual CpG sites and DMRs were then functionally annotated using the ChiPseeker R package^20^. To assess whether these genes were expressed in cord blood, we compared the annotated genes to cord blood RNA-seq datasets from GEO. We further tested for enrichment of FDR-significant CpG site functional location (promoter, exon, intron, intergenic, 5’ UTR, 3’ UTR, and downstream) using a hypergeometric test, to identify locations found in excess or in deficit relative to the respective proportions on the whole array. Following nearest gene assignment of CpGs within FDR-significant regions, canonical pathway analyses were performed using Ingenuity Pathways Analysis (IPA).^21^ To account for the selection criteria of loci/gene coverage on the array and its natural bias towards immune pathways, the IPA Fisher’s exact enrichment tests were done using all genes functionally annotated to CpG sites on the array as a background (i.e. as opposed to the standard analysis which uses all annotated genes in the genome, which would have unnaturally inflated the significance of immune pathways).

Next, a multi-CpG site model of chronological GA was constructed using elastic net penalized linear regression. As input to the model, CpG sites that met an FDR multiple test corrected p-value threshold of 0.05 were included, and the model was adjusted for both infant’s sex and parent-reported race of the infant. EGA estimates were calculated for each sample using leave-one-out cross-validation. In each fold of the cross-validation, the EWAS was reperformed to identify CpG sites with FDR corrected p-values <0.05 that were then used to estimate the model, which was subsequently used to predict the chronological GA of the individual left out of the CpG feature selection and model construction procedure. Using the predicted estimates from the cross-validation in comparison to the chronological GA, the ability of the model to estimate chronological GA was measured using Pearson’s correlation, interclass correlation (ICC), and the percentage of variation explained (ie. linear regression R^2^). We also applied the Bland-Altman method to assess the agreement between chronological GA and EGA^22^.

Finally, association between perinatal environmental factors (obtained by questionnaire) and chronological GA measures were assessed using linear regression. For these analyses, GA was measured in three ways: 1) chronological GA, 2) EGA, and 3) EGAA. EGAA was estimated as the difference between the EGA and the observed chronological GA, such that a positive value would be consistent with biological age acceleration in days. A series of unadjusted (univariate) models with single perinatal environmental exposure were fit for each GA type. To account for the role of multiple environmental factors affecting GA measures, a multi-exposure model was constructed separately for each of the GA measures using a backward selection criterion. In each, the model construction began with the inclusion of all 22 perinatal factors, and the Akaike Information Criteria (AIC) was subsequently utilized to determine those environmental exposures that were retained. The selection was performed using the “step” function in R 4.2.1. statistical programming language^23^.

## Results

### Cohort characteristics

As previously described, WHEALS represents a diverse sampling of the metro-Detroit area^13,14^. The sub-cohort of WHEALS included in this study included 391 individuals, and demographic and pre/peri-natal characteristics for this sample are described in Table 1. Characteristics of our sub-cohort were similar to the full WHEALS cohort, with the exception that our sub-cohort included more married mothers (p=0.011), higher maternal education (4.40×10^−4^), higher chronological GA (p=0.005) and birth weight Z-score (p=007). In terms of race, 62.4% (n=244) of the infant participants had a parent-reported race of African American, 24.3% (n=95) of European American, and 13.3.% (n=52) as other (which included Hispanic, Arabic, and individuals reporting mixed race). The average maternal age at birth was 29.98 years (standard deviation (SD) 5.3), and 56.5% (n=221) of participants lived in an urban setting (defined as having a home in a Detroit-city limit ZIP code) while the remaining 43.5% lived in suburban settings (ZIP code outside of Detroit). 51.4% (n=201) of infants were male, and 63.7% (n=249) of infants were born by vaginal delivery.

### Single CpG site and region-based DNAm associations with chronological GA

Post quality control, 45,296 CpG sites were available for analysis. After correction for multiple tests, 2,435 CpG were associated (FDR-adjusted p-value < 0.05) with chronological GA; the results for these single CpG site analyses are presented in Supplementary Table 1. Of these, 1,330 (55%) were hypermethylated and 1,105 (45%) were hypomethylated, reflecting increased or decreased methylation associated with increasing GA. Overall, 992 unique genes were annotated to these CpGs. To assess whether these genes were expressed in cord blood, we compared the 992 annotated genes to a cord blood RNA-seq dataset from GEO and noted that they are all expressed^24^. The top 40 CpGs that were associated with increased or decreased GA are presented in Supplementary Table 2. The majority of significant CpGs associated with increased GA were located within chromosome 6, with the CpGs annotated to histone genes *H2BC10 and H2BC13*. In comparison, most CpGs associated with decreased GA were spread across different chromosomes.

Next, the CpG sites associated with chronological GA were classified into functional annotation categories within the genome, and the counts/percentages falling into each of these categories are included in Table 2, accompanied by the respective counts/percentages for all of the CpGs included on the array. Comparing the values for each category, there is a clear deviation from a random selection of CpGs associated with chronological GA, with functional annotation categories both significantly over– and under-represented. Specifically, compared to the distribution of all CpGs on the array, enrichment of GA associated CpGs was observed within the first introns (p=3.83×10^−18^) and other introns (p=1.14×10^−9^), 3’ untranslated regions (p=0.016), and promoters located 1-2 kb upstream of the transcription start site (p=0.004). In contrast, there was a significant deficit of GA-associated CpGs within the first 1kb of promoters (p=2.04×10^−48^).

**Table 2.** Functional annotation distribution of CpG sites associated with chronological gestational age. Observed % is the percentage for each annotation compared to what would be expected from CpG sites on the array. Underrepresentation is testing whether we observed fewer than we expected versus overrepresentation is whether we observed more than we expected for each functional annotation, with a p<0.05 being

|  | Observed % | Expected % | P-value |  |
| --- | --- | --- | --- | --- |
|  |  |  | Underrepresentation | Overrepresentation |
| **Promoter (<=1kb)** | 16.3 | 28.7 | $2.04 \times 10^{-48}$ | 1 |
| **Promoter (1-2kb)** | 12.7 | 11 | 0.997 | 0.004 |
| **Promoter (2-3kb)** | 4.9 | 5.2 | 0.271 | 0.760 |
| **5' UTR** | 0.1 | 0.2 | 0.142 | 0.954 |
| **3' UTR** | 2.7 | 2.1 | 0.989 | 0.016 |
| **1st Exon** | 0.5 | 0.5 | 0.460 | 0.658 |
| **Other Exon** | 1.7 | 2.3 | 0.019 | 0.987 |
| **1st Intron** | 17.3 | 11.5 | 1 | $3.83 \times 10^{-18}$ |
| **Other Intron** | 24.5 | 19.7 | 1 | $1.14 \times 10^{-09}$ |
| **Downstream (<=300)** | 0.4 | 0.2 | 0.977 | 0.053 |
| **Distal Intergenic** | 19.0 | 18.7 | 0.674 | 0.345 |

significant.
Footnote: P-value was generated from a hypergeometric test to identify locations found in excess or in deficit relative to the respective proportions on the whole array.

In addition to the single CpG site associations, a region-based analysis was conducted using the *comb-p* approach^18^. In this analysis, 395 regions were significantly associated with GA (FDR adjusted p-values<0.05) (Supplementary Table 3). These regions were comprised of 2,066 CpG sites which were annotated to 336 genes. Of these genes identified by the region-based analysis, 329 (97.9%) overlapped with the 992 identified via the single CpG site analysis, with only seven genes uniquely identified by the region-based approach: *PRSS16, EID3, MIR4706, SCARF2, FBXW2, IKZF5, LYSMD1.* The annotated genes implicated in the region-based analyses were then assessed for canonical biologic pathway enrichment using IPA, using the list of unique genes annotated to CpGs covered on the array as the background for testing. This analysis revealed that the genes near CpGs associated with chronological GA were significantly enriched in pathways related to immune function. Specifically, the top eight most significant pathways (with cutoff p<1×10^−5^) are listed in Figure 1. Enriched pathways included T helper 1 (Th1) and 2 (Th2) activation, macrophage classical activation signaling, and IL-10 signaling. Genes that overlapped with all of the top eight pathways included MHC class II, including *HLA-DMA*, *HLA-DMB, HLA-DQB1*, *HLA-DQB2*, and *HLA-DRB1*. *NFKB1* was enriched in seven out of the top 8 pathways while *CD247*, *IL-6*, and *JAK* were enriched in six out of the top eight. *IL1B* was included in IL-10 signaling and macrophage classical activation signaling, with absence in the T cell enriched pathways.

**Figure 1.**
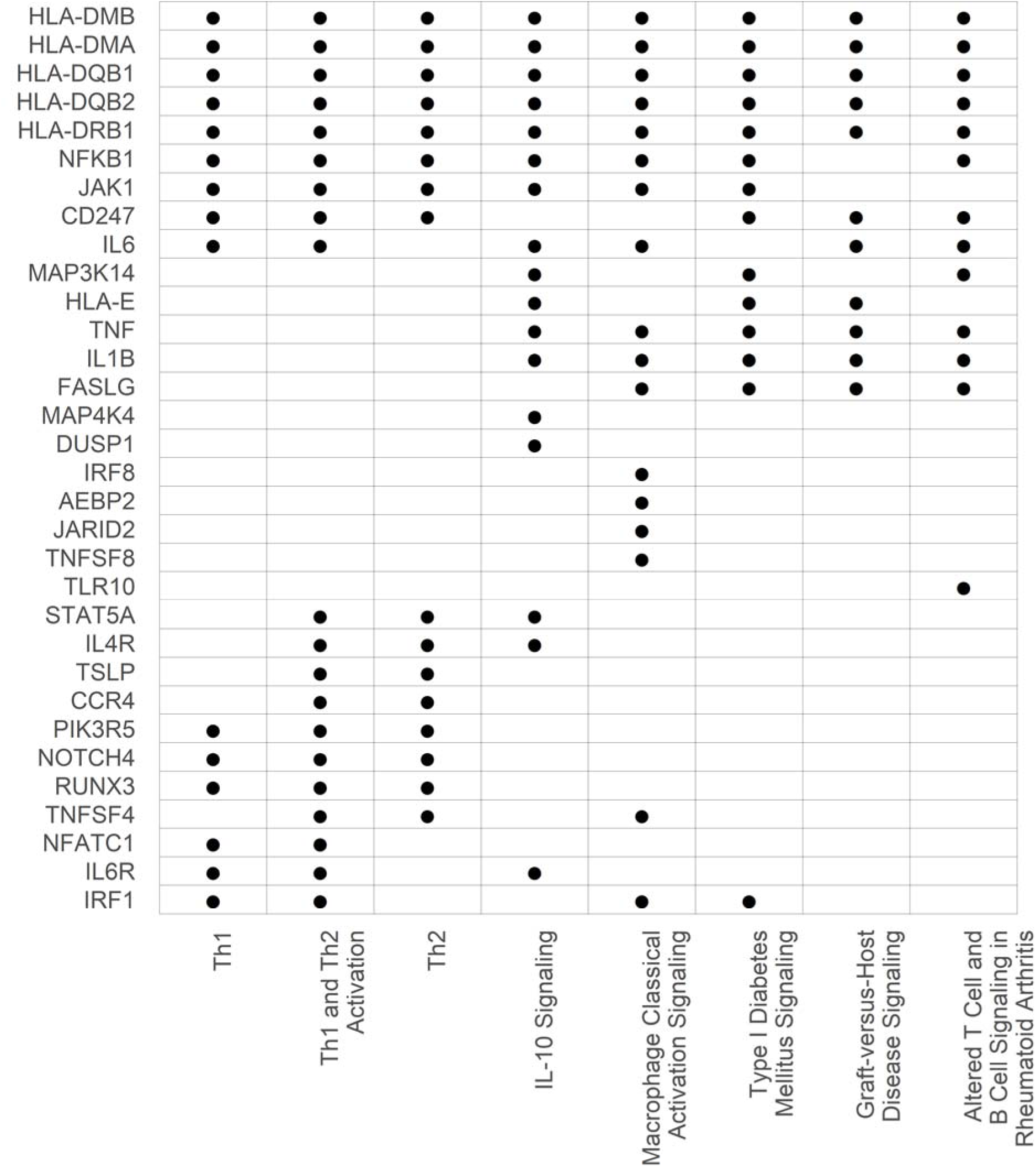
Distribution of genes enriched across top enriched pathways for region-based analyses for chronological gestational age. From the single CpG sites epigenome wide association study with chronological age, we identified differentially methylated regions based on combinations of p-values from the single site EWAS using *comb-p*. These regions were then mapped to genes by ChiPseeker. Pathway analyses via IPA was performed. The table lists the pathways with an enrichment p-value<1×10^−5^. Dot represents enrichment of gene in listed pathway.

### EGA clock

The 2,435 CpG sites associated with chronological GA were utilized to construct a GA clock. The final elastic net penalized linear regression model included 154 CpGs; the regression parameter estimates for this model are included in Supplementary Table 4. Scatter plots of the observed chronological GA versus EGA are shown for the cross-validated and non-cross-validated scenarios in Figures 2a and 2b, respectively. Model assessment metrics demonstrated agreement between the EGA and chronological GA for both cross-validated (Figure 2a; R^2^=0.56, Pearson’s Correlation=0.75, and ICC=0.90) and non-cross-validated (Figure 2b; R^2^=0.86, Pearson’s Correlation=0.93, and ICC=0.90). We also applied the Bland-Altman method as an alternative approach to assess the agreement between chronological GA and EGA, and to assess whether the agreement varied based on preterm, term or post-term GA (Figure 3). The mean difference was calculated at 0.111, with a 95% confidence interval (CI) = – 14.394-14.416. The plot displays strong agreement in the majority of participants, specifically between 35.7-40 weeks. At GA below 35.7 weeks, we see larger negative deviations from the 95% CI between our two measures. To a lesser degree, larger positive deviations are seen at GA>40 weeks.

**Figure 2.**
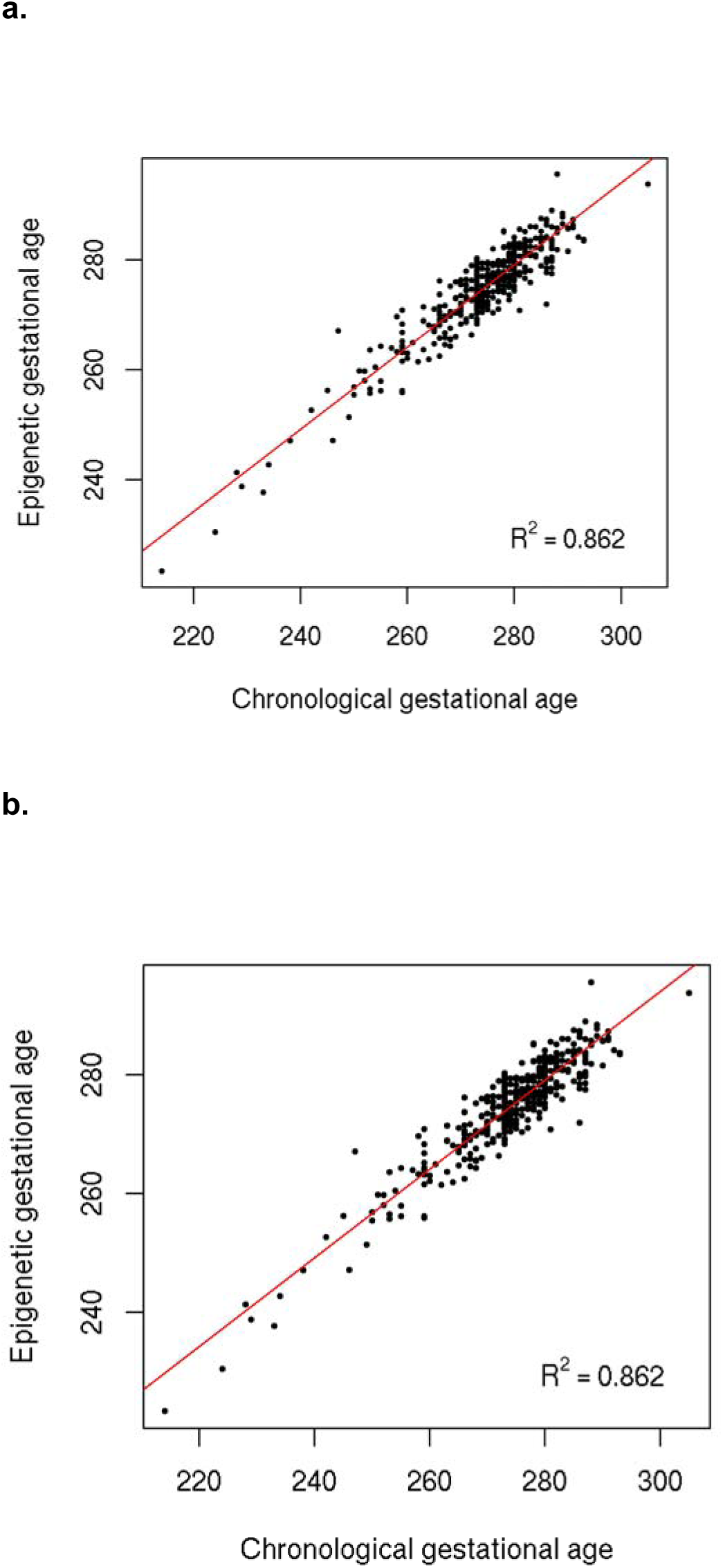
Scatter plots of the observed chronological gestational age (GA) versus calculated epigenetic gestational age (EGA). From the 2,435 CpG sites associated with chronological age, a multi-CpG model was created utilizing elastic net penalized regression model and retained 154 CpGs. The chronological GA was plotted against EGA to assess model assessment agreement. Displayed below are (a) cross-validated and (b) Non-cross validated plots. R^2^ represents the linear regression of the percentage of variation explained.

**Figure 3.**
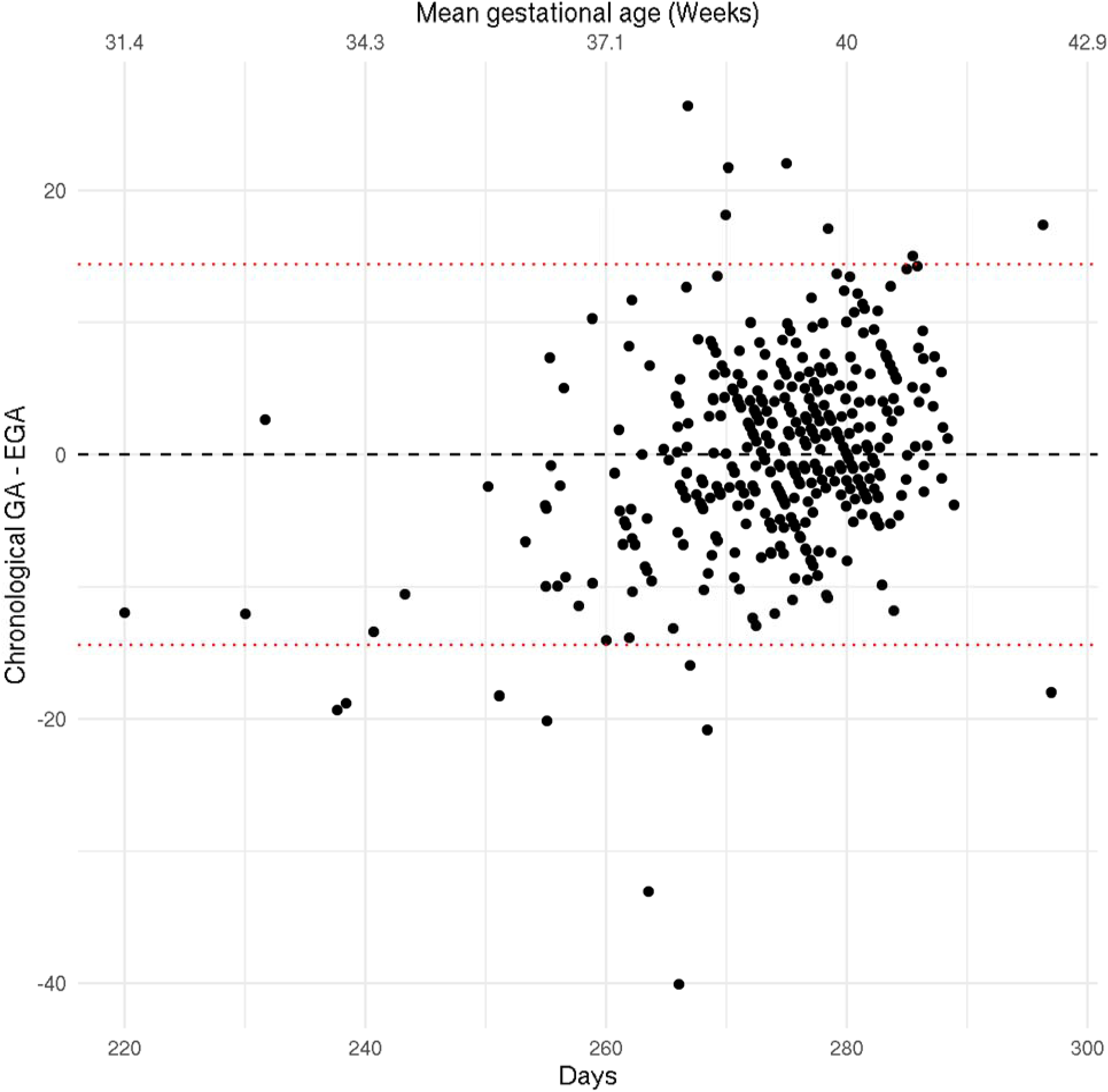
Bland-Altman plot displaying the difference in chronological gestational age (GA) and epigenetic gestational age (EGA) across the mean gestational age in days and weeks. To assess the agreement between chronological GA and EGA across pre-term, term and post-term GA, we applied the Bland-Altman approach below. Red dotted line indicates the confidence interval. The mean difference was calculated at 0.111 and displayed by black dotted line, with a 95% CI = –14.394-14.416. Black dotted line is mean difference.

### Shared biological pathway enrichment between DNAm GA clocks

To compare biological pathways enriched between existing DNAm GA clocks and ours, we performed IPA on the genes mapping to the CpGs comprising each of the four clocks, accounting for the differing backgrounds of genes represented on the respective arrays (Supplementary Table 5-9). The array used for each existing GA clock and number of CpG sites retained in each perspective clock are as follows: Bohlin et al. utilized the Illumina HumanMethylation450^25^ and retained 131 CpGs; Haftron et al. utilized the Illumina MethylationEPIC 850 K^26^ and retained 176 CpGs, and Knight et al. utilized both the Illumina HumanMethylation27 Beadchip and Infinium HumanMethylation450 Beadchip^27^ and retained 148 CpGs. Figure 4 displays the top 10 canonical biological pathways ranked based on the sum of the –log_10_p-values across the four GA clocks. The Knight clock had the most overlapping enriched pathways with the other clocks: ours (n=6), Bohlin (n=6), and Haftron (n=4). The pathway enriched in all clocks was Th1 and Th2 activation, Th2 signaling, protein kinase A signaling, and colorectal cancer metastasis signaling. Six other pathways (molecular mechanisms in cancer, NR1H2 and NR1H3-mediated signaling. RHO GTPase cycle, macrophage classical activation signaling pathway, and tumor microenvironment pathway) were enriched the CpGs from three clocks. Of note, the macrophage classical activation signaling pathway and the Th2 pathway were also captured in our pathway analysis based on all GA-associated CpG regions (Figure 1). Further, the Th1 and Th2 activation pathway enriched in that analysis (Figure 1) was also enriched in our clock CpGs and validated in the Knight clock (Figure 4).

**Figure 4.**
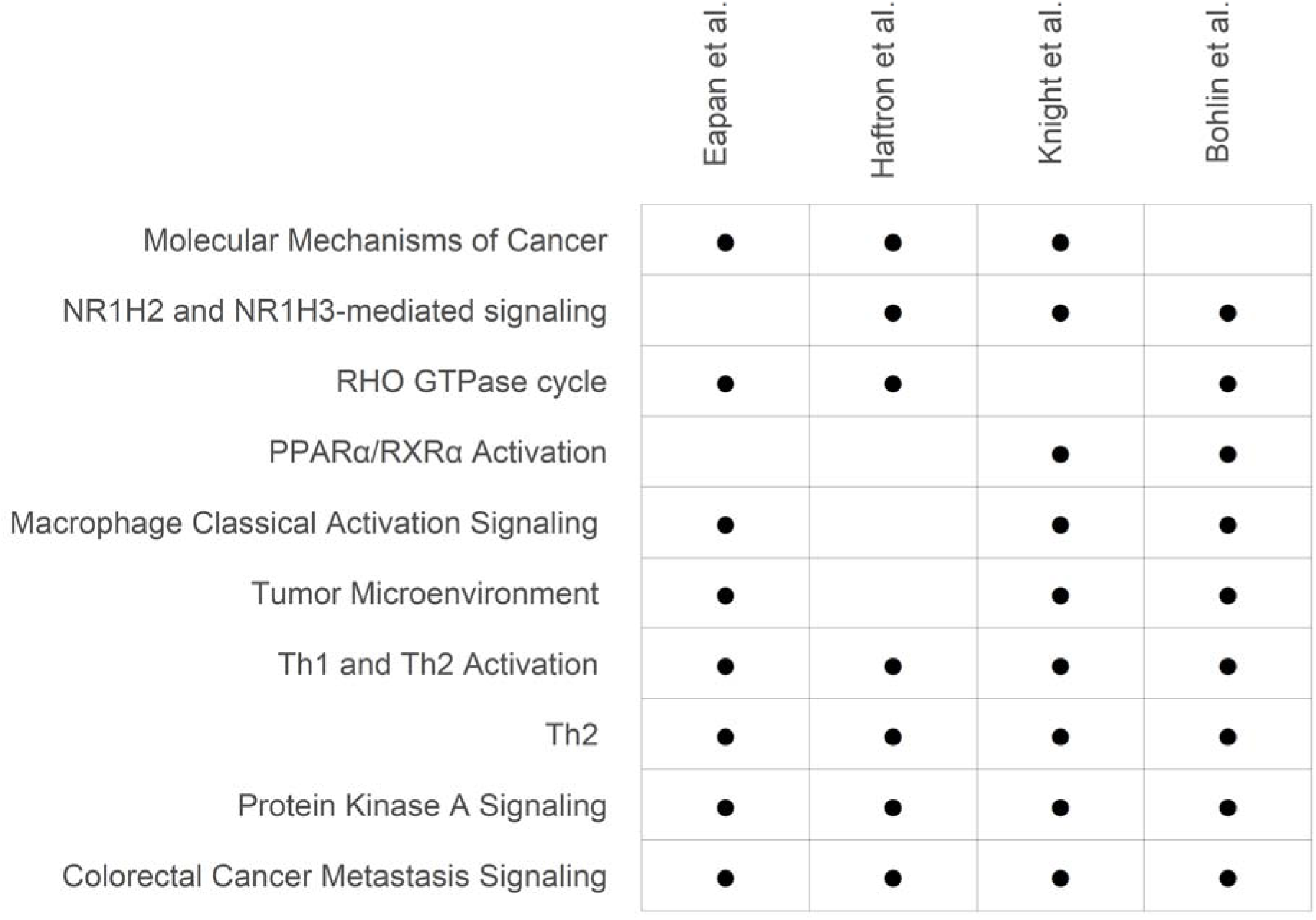
The top 10 biological pathways enriched across gestational age clocks. Pathway analyses via IPA was performed from the genes mapping to the CpGs within each of the four clocks, accounting for the differing background of genes represented on the respective arrays. For each pathway, enrichment p-values were combined across studies using the sum of the study specific pathway enrichment –log10 (p-values) and in this table, the pathways are displayed in their order of significance from top to bottom. Dot represents pathway enrichment p-value<0.05. For all pathways in each study, detailed p-values are listed in Supplementary Table 5.

### Environmental factors and gestational age

Next, individual pre/peri-natal environmental exposures were evaluated for association between chronological GA, EGA, and EGAA in both single and multi-exposure models. The exposures we assessed are included in Table 4. In the single exposure models, there was consistency in direction and magnitude of effects for the exposure that were significantly (p<0.05) associated with chronological GA and EGA, which was expected given the agreement between these two measures shown above (Figure 2). For example, when delivery mode is stratified by vaginal, planned and unplanned C-section, planned C-section was significantly associated with a lower chronological GA and EGA, while unplanned C-section was associated with higher chronological GA (Table 3). Compared to children of mothers without a previous pregnancy, children of mothers with ≥4 previous pregnancies had both lower chronological GA (Coefficient [95% CI] = –3.67 [-7.34,-0.01], p=0.051) and lower EGA (Coefficient [95% CI] = –4.15 [-7.05, –1.26], p=0.005). Birth weight z-score was associated with higher chronological GA and EGA (Coefficient [95% CI] = 2.40 [1.28, 3.51], p<0.001, 2.05 [1.17, 2.94], p<0.001)(Table 3). Unplanned C-section was the sole exposure significantly associated with EGAA, with its negative direction reflecting a decrease in EGAA (Coefficient [95% CI] = –2.08 days [-4.07, –0.10], p=0.040), relative to vaginal delivery.

**Table 3.** Single pre-and peri-natal exposure associations with gestational age measures. For each gestational age (GA) measure, unadjusted univariate linear regression models with single prenatal environmental exposures were fit for each GA type.

| Covariate | Levels | Chronological Gestational Age |  |  | Epigenetic Gestational Age |  |  | Epigenetic Gestational Age Acceleration |  |  |
| --- | --- | --- | --- | --- | --- | --- | --- | --- | --- | --- |
|  |  | Coefficient | 95% CI | *P-value | Coefficient | 95% CI | *P-value | Coefficient | 95% CI | *P-value |
| Maternal Age |  | 0.01 | (-0.19, 0.22) | 0.891 | -0.08 | (-0.24, 0.09) | 0.368 | -0.09 | (-0.23, 0.05) | 0.203 |
| Paternal Age |  | -0.10 | (-0.27, 0.07) | 0.237 | -0.13 | (-0.26, 0.01) | 0.064 | -0.03 | (-0.14, 0.08) | 0.648 |
| BMI First Recorded in Pregnancy |  | -0.05 | (-0.18, 0.09) | 0.490 | -0.07 | (-0.18, 0.04) | 0.198 | -0.02 | (-0.11, 0.07) | 0.621 |
| BMI Last Recorded in Pregnancy |  | 0.02 | (-0.12, 0.16) | 0.775 | -0.01 | (-0.12, 0.10) | 0.851 | -0.03 | (-0.13, 0.06) | 0.524 |
| Birth Weight z-score |  | 2.40 | (1.28, 3.51) | <0.001 | 2.05 | (1.17, 2.94) | <0.001 | -0.35 | (-1.11, 0.42) | 0.372 |
| Parity |  | -0.42 | (-1.40, 0.55) | 0.393 | -0.89 | (-1.66, -0.13) | 0.022 | -0.47 | (-1.12, 0.18) | 0.155 |
| Marital Status | Married | 2.08 | (-0.24, 4.40) | 0.078 | 1.41 | (-0.43, 3.24) | 0.134 | -0.68 | (-2.23, 0.87) | 0.392 |
| Urban Residence | Urban (Detroit) | -0.98 | (-3.19, 1.23) | 0.382 | -1.27 | (-3.02, 0.48) | 0.154 | -0.28 | (-1.76, 1.19) | 0.705 |
| Household Income (ref = <20k) | 20k-<40k | 2.76 | (-1.07, 6.58) | 0.157 | 2.70 | (-0.33, 5.74) | 0.081 | -0.05 | (-2.63, 2.52) | 0.967 |
|  | 40k-<80k | 3.86 | (0.22, 7.49) | 0.038 | 2.93 | (0.04, 5.81) | 0.047 | -0.93 | (-3.38, 1.51) | 0.454 |
|  | 80k-<100k | 2.02 | (-2.24, 6.28) | 0.351 | 1.32 | (-2.06, 4.71) | 0.442 | -0.70 | (-3.56, 2.17) | 0.632 |
|  | >=100k | 6.28 | (2.08, 10.48) | 0.003 | 4.51 | (1.17, 7.84) | 0.008 | -1.78 | (-4.60, 1.05) | 0.217 |
|  | refuse to answer | 1.84 | (-2.47, 6.14) | 0.402 | 2.61 | (-0.81, 6.02) | 0.135 | 0.77 | (-2.13, 3.66) | 0.603 |
| Maternal Education (ref = <= HS diploma) | HS diploma | -4.14 | (-10.23, 1.94) | 0.182 | -1.71 | (-6.55, 3.14) | 0.489 | 2.43 | (-1.67, 6.53) | 0.244 |
|  | Some college | -5.39 | (-10.98, 0.19) | 0.058 | -2.94 | (-7.38, 1.51) | 0.195 | 2.46 | (-1.30, 6.22) | 0.200 |
|  | Bachelor's degree | -1.57 | (-7.23, 4.09) | 0.586 | -0.44 | (-4.95, 4.07) | 0.848 | 1.13 | (-2.68, 4.95) | 0.560 |
| Delivery Mode (ref = vaginal) | Planned C-section | -4.27 | (-7.08, -1.45) | 0.003 | -4.31 | (-6.53, -2.08) | <0.001 | -0.04 | (-1.95, 1.87) | 0.966 |
|  | Unplanned C-section | 3.41 | (0.48, 6.33) | 0.022 | 1.32 | (-0.99, 3.64) | 0.262 | -2.08 | (-4.07, -0.1) | 0.040 |
| First Born | Yes | -0.50 | (-2.81, 1.80) | 0.667 | 0.85 | (-0.97, 2.67) | 0.360 | 1.35 | (-0.18, 2.88) | 0.083 |
| Number of Previous Pregnancies (ref = 0) | 1 | 0.19 | (-3.01, 3.40) | 0.905 | -0.23 | (-2.75, 2.29) | 0.858 | -0.42 | (-2.56, 1.72) | 0.698 |
|  | 2 | -0.89 | (-4.03, 2.26) | 0.580 | -2.36 | (-4.84, 0.11) | 0.061 | -1.48 | (-3.58, 0.63) | 0.168 |
|  | 3 | -2.56 | (-6.28, 1.16) | 0.177 | -1.53 | (-4.46, 1.40) | 0.304 | 1.03 | (-1.46, 3.51) | 0.417 |
|  | >=4 | -3.67 | (-7.34, 0.01) | 0.051 | -4.15 | (-7.05, -1.26) | 0.005 | -0.49 | (-2.94, 1.97) | 0.698 |
| Infant Gender | female | 0.62 | (-1.58, 2.81) | 0.581 | 0.77 | (-0.97, 2.51) | 0.384 | 0.15 | (-1.31, 1.62) | 0.836 |
| Maternal Antibiotic Use during Pregnancy | Yes | -0.46 | (-2.69, 1.77) | 0.683 | -0.03 | (-1.79, 1.73) | 0.974 | 0.43 | (-1.08, 1.95) | 0.573 |
| Maternal Antifungal Use during Pregnancy | Yes | -0.81 | (-3.71, 2.10) | 0.586 | -1.01 | (-3.31, 1.29) | 0.388 | -0.20 | (-2.18, 1.77) | 0.840 |
| Season of Birth (ref = winter) | Spring | -1.08 | (-4.44, 2.28) | 0.529 | -1.26 | (-3.92, 1.40) | 0.352 | -0.19 | (-2.42, 2.05) | 0.870 |
|  | Summer | -1.70 | (-4.86, 1.46) | 0.292 | -0.46 | (-2.96, 2.05) | 0.721 | 1.24 | (-0.87, 3.35) | 0.248 |
|  | Fall | -1.42 | (-4.57, 1.73) | 0.376 | -0.68 | (-3.18, 1.81) | 0.591 | 0.74 | (-1.36, 2.84) | 0.489 |
| Environmental Tobacco Smoke | Yes | -1.70 | (-4.22, 0.81) | 0.184 | -1.39 | (-3.38, 0.60) | 0.171 | 0.32 | (-1.37, 2.00) | 0.712 |
| Mom Smoking Status | Yes | -2.23 | (-5.69, 1.24) | 0.207 | -2.44 | (-5.18, 0.30) | 0.081 | -0.21 | (-2.53, 2.10) | 0.856 |
| Indoor Pets | Yes | 0.85 | (-1.42, 3.12) | 0.464 | -0.16 | (-1.96, 1.64) | 0.862 | -1.01 | (-2.52, 0.51) | 0.192 |
| Outdoor Pets | Yes | 2.26 | (-3.27, 7.79) | 0.422 | 1.14 | (-3.24, 5.53) | 0.608 | -1.12 | (-4.81, 2.58) | 0.553 |
| Maternal Race (ref = African American) | European American | 1.36 | (-1.27, 3.99) | 0.311 | 1.49 | (-0.59, 3.57) | 0.161 | 0.13 | (-1.63, 1.89) | 0.885 |
|  | Other | 1.27 | (-1.93, 4.46) | 0.437 | 0.98 | (-1.55, 3.51) | 0.445 | -0.28 | (-2.42, 1.85) | 0.794 |
| Infant's Race (ref = African American) | European American | 1.50 | (-1.12, 4.12) | 0.261 | 1.54 | (-0.53, 3.62) | 0.144 | 0.05 | (-1.71, 1.80) | 0.960 |
|  | Other | 2.19 | (-1.11, 5.50) | 0.193 | 1.40 | (-1.21, 4.02) | 0.292 | -0.79 | (-3.00, 1.42) | 0.483 |

Abbreviations: CI, confidence interval, C-section, Caesarean section, GA, gestational age; HS, high school; k, thousand; ref, reference.
Coefficient: Regression parameter estimate for CpG site - gestational age model
\*F-test evaluating significant differences for continuous or dichotomous characteristics for each GA outcome

**Table 4:**
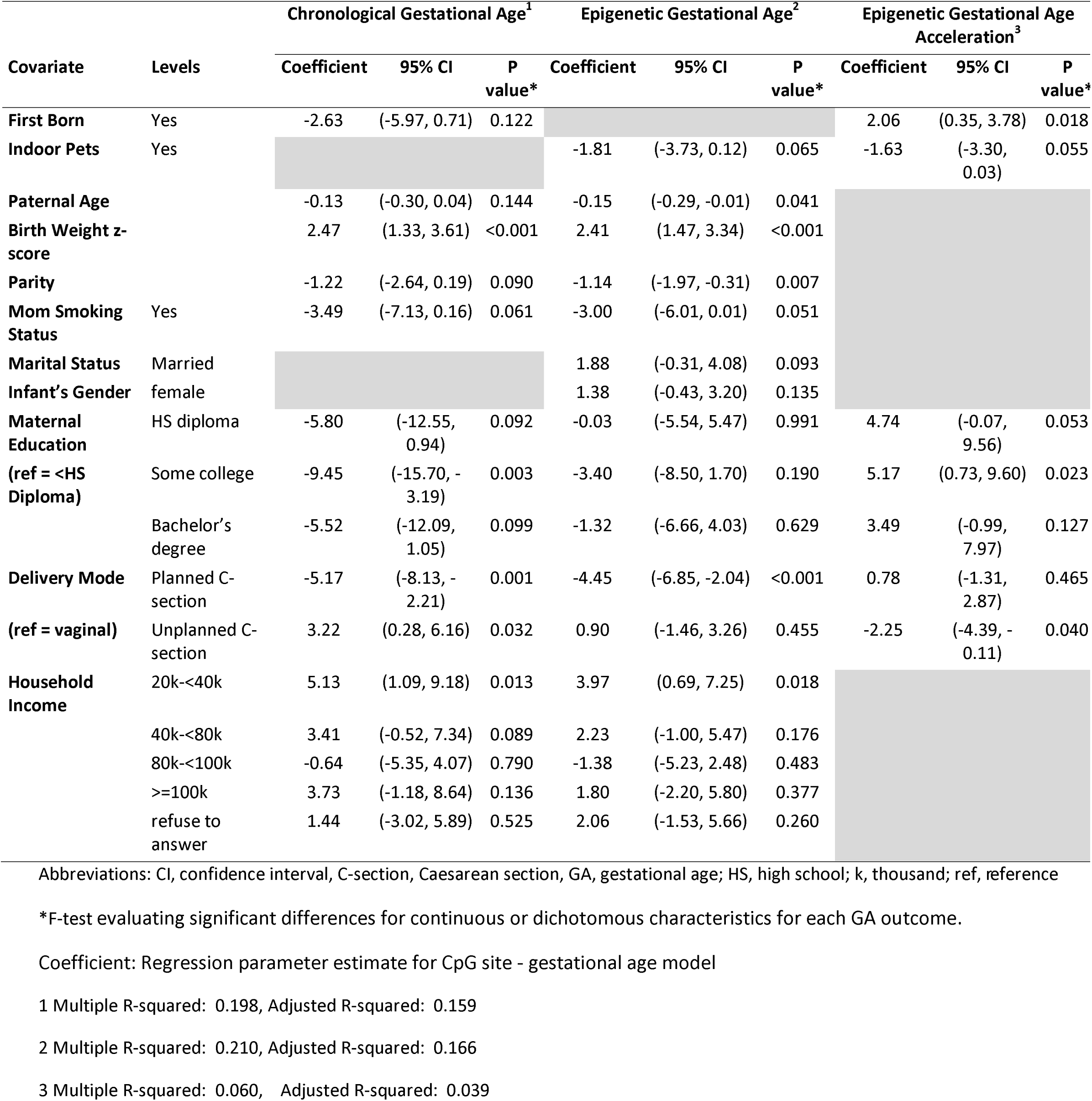
Summary of multivariable model selected by backward stepwise selection. To account for the role of multiple environmental factors affecting gestational age (GA) measures, a multi-exposure model was constructed separately for each GA measure using a backward selection criterion. In each, the model construction began with the inclusion of all 22 prenatal factors (those listed in Table 3 excluding infant’s race), and the Akaike Information Criteria (AIC) was utilized to determine those environmental exposures that were retained.

To assess whether groups of environmental factors were associated with GA assessments, a multi-exposure model was constructed for each of the three GA measures using a backward selection procedure, and the resulting multi-exposure models are summarized in Table 4. The pre/perinatal multi-exposure models explained more of the variation in chronological GA and EGA (adjusted R^2^ = 0.159 and 0.166, respectively) in comparison to EGAA (adjusted R^2^ = 0.039). Further, there was general agreement between multi-exposure models of both chronological GA and EGA, and the directions of effect also agreed with their single exposure estimates. Environmental factors retained in models for both chronological GA and EGA were paternal age, birth weight, parity, maternal smoking status, maternal education, delivery mode, and household income. Indoor pets was retained in the model for EGA, but not chronological GA, while first born status was retained in the model for chronological GA but not EGA. In the EGA and chronological GA models, birth weight, parity, household income, and mode of delivery were statistically significant (p<0.05). In the multi-exposure model of EGAA, prenatal indoor pets, maternal education level, delivery mode, and firstborn status were retained. However, within this model, only unplanned C-section, maternal education, and firstborn status were statistically significant (p<0.05). Among these, unplanned C-section was associated with decreased EGAA, while first born status and maternal education (some college) were associated with increased EGAA.

## Discussion

In our study of DNAm and GA, our region-based EWAS results identified the enrichment of Th1 and Th2 signaling, IL10 signaling, and macrophage classical activation being significantly enriched in DNAm associated with chronological GA. Additionally, CpG sites retained in our GA clock also highlight these Th1, Th2 and IL-10 signaling pathways. The early life immune system is noted to be associated with GA, with highlighted pathways involving pro– and anti-inflammatory immune mechanisms and variations in T cell populations^28^.

In our single CpG site and region-based EWAS, HLA-class II genes (*HLA-DMA, HLA-DMB, HLA-DQB1, HLA-DQB2, and HLA-DRB1*) and *NFKB1* were the most significant genes implicated in pathways significantly associated with chronological GA. Among the HLA-class II genes identified, expression of HLA-DRB1 has been previously reported to be associated with both age-related changes in brain structure and cognitive performance^29,30^ and also with longevity beyond 85 years of age. Further, single cell proteomic studies of cord blood have found associations of GA with both T-reg pathways and NFKB1 signaling in antigen presenting cells expressing HLA-class II genes^28^, and T-cell pathway skewing and chronic antigen stimulation have been previously noted to mediate immunosenescence^31^. Although all components of the innate and adaptive immune system are adversely affected to varying extents by aging, antigen presentation, T cell activation, and NFKB1 signaling appear to be particularly sensitive and important to the aging process^32^. Our findings highlight the role of DNAm at HLA-class II genes and *NFKB1* that may be regulating these immune pathways through epigenetic gene regulation.

We herein present a GA clock based on the Asthma&Allergy array that is correlated with chronological GA (r = 0.75). As DNAm arrays have evolved, the accuracy of GA prediction has also changed. Our clock had a stronger correlation than some but not all previously reported GA clocks: Bohlin clock (r=0.61)^25^, Knight clock (r=0.91)^27^, and most recently the Haftron clock (r=0.72)^26^. This variation in correlation between chronological GA and EGA is likely due to the differences in both the number and epigenome-wide coverage of CpGs. However, our findings suggest that the accuracy of our DNAm-based GA clock is comparable to other similar existing methods particularly between 35-40 weeks.

When comparing previously published GA clocks to ours based on enriched pathways, there were multiple findings of note. First, three of the immune related pathways (macrophage classical activation signaling pathway, Th2 pathway, and Th1 and Th2 activation pathway) captured in our pathway analyses based on 1) all GA-associated region CpGs and 2) those CpGs in our clock were found to be enriched in at least one of the other three clocks. These findings not only validate the importance of immune related pathways in early life aging, which is consistent with the effects demonstrated in the broader aging literature^33^, but they also reveal that these immune pathways are also important in the prediction of GA. Second, these analyses across the four clocks implicate multiple cancer associated pathways. This finding is consistent with the literature implicating accelerated biological aging as associated with incidence across multiple tumor types^34^.

The environment is noted to have effects on DNAm, and consistently, our findings support the impact of multiple pre/peri-natal factors on GA and EGAA. Previous studies investigating the association between perinatal environmental factors and DNAm with GA have primarily focused on maternal depression, prenatal medication, and smoking^12^. Our study has taken a more agnostic approach, investigating a longer list of pre/peri-natal environmental factors. Further, to our knowledge, there have been no previously published studies of pre/peri-natal exposures associated with EGAA.

While maternal smoking has been associated with placental DNAm and decreased GA^35^, our findings did not find a significant association of maternal smoking with chronological GA, EGA, or EGAA. SES, specifically low SES, has been associated with decreased GA^36^. Our findings show the reverse for those with household incomes less than $20,000, but these findings were also similar for those with household incomes greater than $100,000, both associated with increased chronological GA and EGA. Parity has shown a correlation with chronological GA and preterm births^37^. Our findings show that it is also associated with decreased EGA. Lastly, unplanned C-section but not planned C-section affected EGAA, potentially due to clinically-induced activation of labor that has different biological effects than those from naturally occurring pathways activating birth.

Our analyses of environmental variables also emphasize the impact of multiple pre-and perinatal environmental factors in chronological GA, EGA and EGAA. Specifically, in our backward selection model fit, prenatal indoor pets was retained in the multi-exposure model for EGA and EGAA, with prenatal indoor pets associated with lower EGA and decreased EGAA. We and others have shown the protective effect of prenatal pet exposure on allergic outcomes such as asthma and atopic dermatitis^38–40^. Further, other studies have found that EGAA is a risk factor for multiple childhood onset conditions^7,8^. While more studies are needed to evaluate whether EGAA mediates the effect of prenatal pet exposure on risk of these childhood onset outcomes, including allergic conditions, we wanted to highlight this possibility to incent further investigation of potential contributions of multiple environmental exposures to EGAA (and biological age acceleration in general) and risk of disease.

There are several strengths to our study. These include, but are not limited to, early DNAm assessment across preterm, term and post-term GA, a large diverse cohort (in terms of race and SES), and detailed perinatal environmental exposure data. However, limitations to our study also exist. One such limitation is that our characterization of C-section as planned versus unplanned was chart abstracted and defined as whether the C-section was scheduled or not.

Although most of the unplanned C-sections were likely laboring, there may be a small portion that were not (e.g. maternal preeclampsia, large-for-gestational age fetus). Future studies further stratifying delivery mode by labor status would help clarify the effects of labor on EGA and EGAA. Further, while we evaluated multiple exposure associations with GA measures, the list was not exhaustive. Given our suggestive findings, future studies should expand the list of exposures investigated, including perinatal environmental exposures such as indoor/outdoor environmental pollution measures which have been shown to impact DNAm^41,42^.

In conclusion, our findings highlight immune pathway and gene associations with cord-blood EGA. Our findings additionally show influence of perinatal factors on chronological GA, EGA and EGAA, which may have an influence on subsequent biological pathways throughout life. Future studies applying EGA clocks to disease outcomes would benefit from incorporating the influence of perinatal environmental factors to identify mechanisms in risk of disease.

## Data sharing

Datasets analyzed in this study are not publicly available; consent for public release of epigenetic data from participants was not obtained from participants. However, data to generate figures and tables are available from the corresponding author with the appropriate permission from the WHEALS study team and investigators upon reasonable request and Institutional Review Board approval.

## Author contributions

AAE: Formal analysis, funding acquisition, investigation, methodology, project administration, data curation, conceptualization, resources, supervision, writing – original draft, review, edit, visualization, and validation.

IL: Formal analysis, methodology, writing – original draft, review, edit, data curation, investigation.

MP: Formal analysis, methodology, writing – original draft, review, edit, data curation, investigation.

XL: Writing – review, edit, analysis and interpretation

AU: Writing – review, edit, project administration

JS: Writing – review, edit, analysis and interpretation

ACB: Writing – review, edit, analysis and interpretation

AS: Writing – review, edit, analysis and interpretation

NS: Writing – review, edit, interpretation

ET: Writing – review, edit, methodology, project administration, formal analysis

LK: Writing – review, edit, analysis and interpretation

CO: Writing – review, edit, methodology, study design

CJ: Conceptualization, data curation, funding acquisition, investigation, methodology, project administration, supervision, writing – review, edit

EZ: Writing – review, edit, study design, conceptualization

AL: Formal analysis, funding acquisition, investigation, methodology, project administration, data curation, conceptualization, resources, supervision, writing – original draft, review, edit, visualization, validation.

All authors have reviewed and edited the article before submission, agreed on the journal before submission and agree to take responsibility and be accountable for its contents.

## Abbreviations

DNAm: DNA methylation
GA: gestational age
EGA: epigenetic gestational age
EGAA: epigenetic gestational age acceleration
EWAS: epigenome-wide association study
Th1: T helper 1
Th2: T helper 2
WHEALS: Wayne County Health, Environment, Allergy and Asthma Longitudinal Study

